# Amino acids bind to phase-separating proteins and modulate biomolecular condensate stability and dynamics

**DOI:** 10.1101/2025.03.05.641737

**Authors:** Xufeng Xu, Merlijn H. I. van Haren, Iris B. A. Smokers, Brent S. Visser, Paul B. White, Robert S. Jansen, Evan Spruijt

## Abstract

Biomolecular condensates (BCs) are versatile intracellular compartments that help organize intracellular space and perform diverse biological functions. In living cells, BCs are surrounded by a complex environment, of which amino acids (AAs) are prominent components. However, it is unclear how AAs interact with condensate components and influence the formation and material properties of condensates. Here, we show that phase separation is suppressed with increasing glycine concentration by using model heterotypic condensates made of nucleophosmin 1 (NPM1) and ribosomal ribonucleic acid (RNA): the condensate density is decreased and the dynamics inside the condensate are increased. We find that glycine weakly binds to amide groups in the protein backbone and aromatic groups in the side chains with a 1∼2 M affinity, weakening the backbone-backbone interactions between neutral and charged disordered proteins while strengthening the interactions between aromatic stickers. This leads to different modulations of the phase behaviour in condensates formed by cation/π-π interactions and condensates formed by charge complexation. We further show that a similar modulation effect on BCs is observed for other proteinogenic AAs and can be transferred to short peptides. These insights offer a better understanding of BC stability, with implications for new strategies to modulate the dynamic properties of BCs *in vivo*.

## Introduction

Biomolecular condensates (BCs) are condensed bodies in the cell, which normally form through liquid-liquid phase separation^1^. They are composed of a rich variety of proteins and nucleic acids^2^, and are found in a wide range of cell types and developmental stages^3^. The largest and earliest observed BC is the nucleolus, which was reported in the 1830s^4^. Only recently, the formation process and physical properties of these BCs were studied more systematically by Brangwynne, Hyman, Rosen, and others^5–7^. Subsequently, the fundamental roles of BCs in cellular homeostasis and disease have been extensively studied^8,9^. The phase-separated nature of BCs makes them sensitive to various types of changes in their environment, including crowding, pH, ionic strength, and the presence of modulators in the form of ATP, peptides, and other guest molecules^10–12^.

Amino acids (AAs) are known to constitute a major component of the intracellular milieu. Strikingly, more than 25% of the total volume and 6% of the total dry mass of a mammalian cell was reported to be taken up by free AAs^13^. It has been recently found that AAs have a general effect on modulating protein-protein interactions^14,15^ and some AAs can modulate the formation process of stress granules^16^. It has also been shown that specific AAs can impact the formation of BCs. For example, Paccione et al.^17^ reported that glutamate (E) enhanced the condensate formation of bacterial cell division protein FtsZ and its DNA-bound regulator SlmA. Kozlov et al.^18^ also reported that glutamate promoted the formation of condensates of *E. coli* SSB protein. However, it remains unclear how AAs interact with components that form BCs and how the material properties of BCs are modulated as a result of these interactions.

To establish if AAs could impact BC formation, we studied the effect of glycine on the phase separation of nucleophosmin (NPM1) and ribosomal ribonucleic acid (RNA), which is an *in vitro* heterotypic condensate model of nucleoli^19^. We find that the miscibility gap decreases, while protein dynamics inside the condensates increase with the addition of glycine. Interestingly, except for glutamate (E), all tested amino acids have a similar effect on NPM1/RNA condensates as glycine, suggesting that an interaction between the AA backbone and the condensate components underlies the general modulation effect.

To elucidate the mechanism behind the modulation effect, we studied the effect of glycine on four different synthetic condensate systems with different intermolecular interactions driving condensation: K_72_/ATP^20^, polyLys/polyAsp (K_10_/D_10_)^21^ (both formed by electrostatic interaction), FFssFF^22^ (formed by π-π stacking) and WGR-4 peptide^23^ (formed mainly by cation-π interactions). By the combinative use of nuclear magnetic resonance (NMR), liquid chromatography-mass spectrometry (LC-MS), and microplate reader assays, we find that AAs bind weakly to amide groups in the protein/peptide backbone and aromatic groups^24^ in the side chain (*K*_d_ ≈ 1∼2 M), primarily through AAs’ amine group in the main chain. This leads to the partitioning of AAs inside BCs (*K*_p_ ≈ 5). We hypothesize that the binding of AAs increases the effective dielectric permittivity in the vicinity/microenvironment around the binding sites^25–27^. This leads to the weakening of protein backbone-backbone interactions, which have so far not received much attention in the context of condensate formation, and the strengthening of cation/π and π-π interactions. Overall, AAs can thus enhance condensation driven predominantly by cation/π-π interactions, and suppress condensation driven by charge complexation.

Interestingly, we also find that short peptides with up to 8 amino acids exhibit similar effects on NPM1/rRNA condensates at the same AA residue concentration, suggesting that they can bind to protein backbones in an additive manner. These findings open up a new molecular design platform to fine-tune the material properties of BCs, which may find applications in regulating protein functions *in vivo* and treating condensate-related diseases.

## Results and discussion

### Glycine modulates the stability and material properties of NPM1-RNA condensates

We use a heterotypic condensate model of the granular component of the nucleolus, consisting of NPM1 and ribosomal RNA^28^. NPM1 and RNA undergo liquid-liquid phase separation under physiological conditions (10 mM Tris, 150 mM NaCl, pH 7.5), leading to the formation of well-defined spherical condensates, enriched in both NPM1 and RNA (**Figure 1a**) with a substantial NPM1 miscibility gap (width of the two-phase region), which agrees with previous findings^19,29^. To investigate if amino acids (AAs) impact NPM1/RNA phase separation, we selected glycine (G) as the simplest AA to minimize additional effects from AA side chains. With increasing concentrations of glycine, NPM1/RNA condensates became gradually less bright and less spherical, suggesting that their local density and surface tension decreased (**Figure 1b**) . We quantified the concentrations of NPM1 in the dilute and condensed phase as a function of glycine concentration using fluorescence spectroscopy and microscopy (experimental details in **Methods**) and found that the NPM1 concentration in the dilute phase increased by ∼50% from ∼5.9 µM up to ∼8.6 µM, while the concentration in the condensate decreased 6-fold from ∼228 µM down to ∼36 µM, reaching a plateau at around 0.6 M glycine (**Figure 1c** and **1d**). However, the partitioning of RNA was affected only slightly (**Figure S1**), suggesting the glycine weakened NPM1-NPM1 or NPM1-RNA interactions, while RNA-RNA interactions remained unchanged. We also observed a similar effect of glycine on NPM1 condensates formed in the presence of the crowding agent PEG (10k Da)^29^ (**Figure S2**).

**Figure 1:**
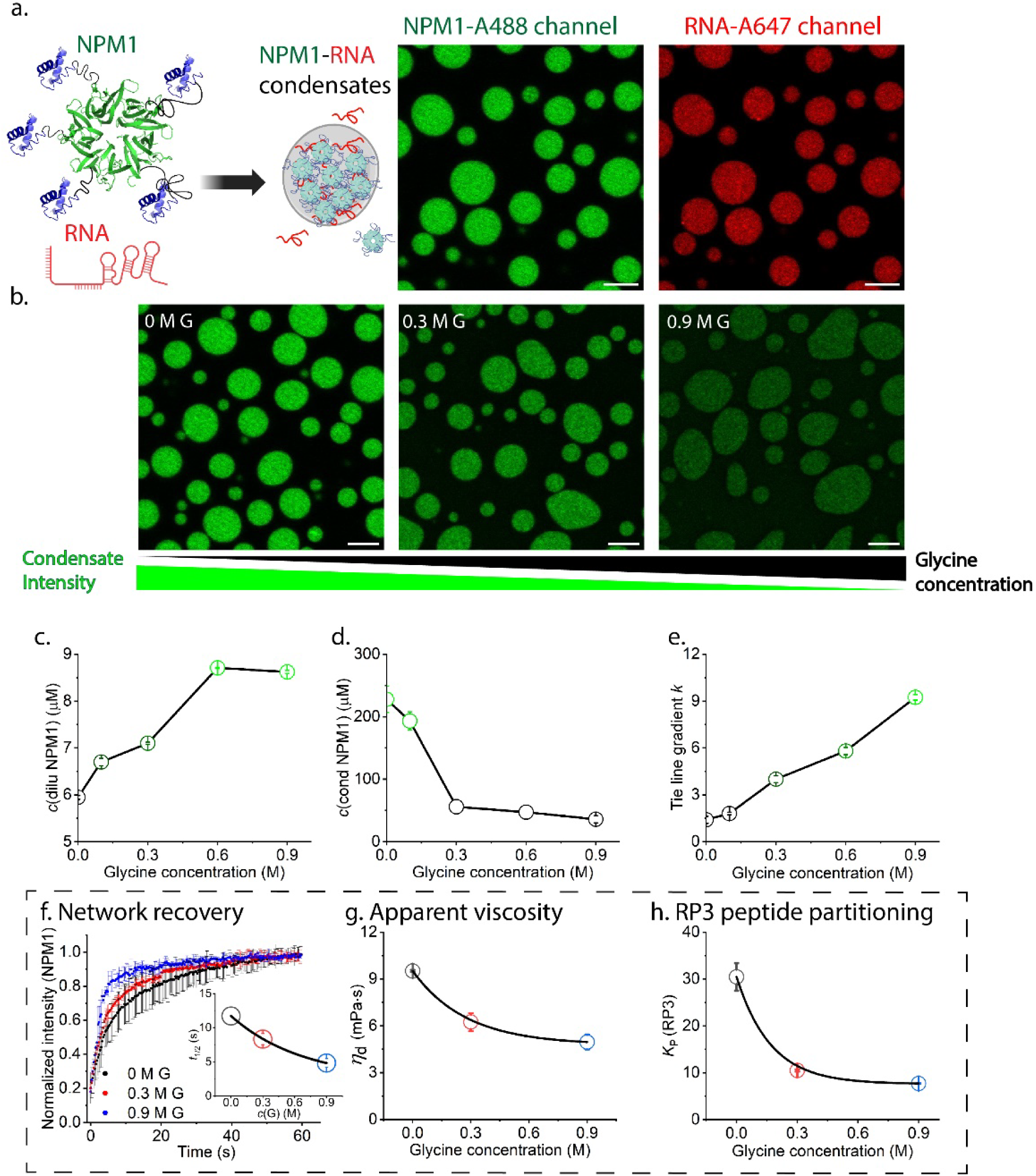
The phase behaviour and material properties of NPM1-RNA condensate after the addition of glycine. a. Schematic illustration of NPM1 protein structures (oligomerization domain (green, PDB: 4N8M) connected via disordered regions (grey) to the C-terminal nucleic acid binding domain (blue, PDB: 2VXD) and their formation of condensates with RNA. Fluorescence confocal microscopy images of NPM1-RNA condensates in NPM1-Alexa488 channel and RNA-Alexa647 channel; Scale bar = 5 μm. b. Confocal fluorescence microscopy images of NPM1-RNA condensates in NPM1-Alexa488 channel after the addition of 0, 0.3, and 0.9 M glycine; c and d. NPM1 concentrations in the dilute and condensate phases after the addition of glycine (0, 0.1, 0.3, 0.6, and 0.9 M) (calculation details in **Methods**); e. Calculated tie line gradient *k* after the addition of glycine (0, 0.1, 0.3, 0.6, and 0.9 M); f. Average FRAP recovery curves of NPM1 and the calculated recovery half-life (*t*_1/2_) after the addition of glycine (0, 0.3, and 0.9 M); g. Apparent viscosity (*η*_d_) of fluorescein (Alexa Fluor 488) in NPM1-RNA condensates after the addition of glycine (0, 0.3, and 0.9 M); h. Partitioning coefficients (*K*_p_) of RP3 in NPM1-RNA condensates after the addition of glycine (0, 0.3, and 0.9 M).

To further quantify the effect of glycine on intermolecular interactions underlying phase separation, we calculated the tie-line gradient *k,* according to Qian et al^30^, which can be expressed by effective interaction difference for the condensate formation 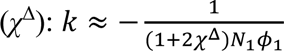where *ϕ*_1_ and *N*_1_ denote the volume fraction and length of the component 1 (NPM1), respectively, and 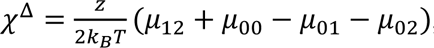, with *µ* the contact energy between the solvent (0), solute 1 (NPM1) and 2 (RNA), respectively, *z* a coordination constant, *k*_B_ Boltzmann’s constant and *T* the absolute temperature^30^. As shown in **Figure 1e**, the tie-line gradient *k* increases with increasing glycine concentration, indicating a weaker associative interaction driving the condensate formation.

The weaker associative interaction in the condensate phase (i.e. decreased condensate stability) in turn modulates the local dynamic properties of the condensates. As shown in **Figure 1f**, NPM1 recovers faster after photobleaching at high glycine concentration. The calculated recovery half-life (*t*_1/2_) for NPM1 in the condensates decreased by more than a factor of two (from 12 s to 5 s) after the addition of 0.9 M glycine. By contrast, the recovery half-life of RNA hardly changed with the addition of glycine (**Figure S3**). In agreement with the FRAP experiment, the effective viscosity of the condensate, as estimated from the diffusion of free fluorescein (Alexa Fluor 488) molecules by raster image correlation spectroscopy (RICS), decreased by a factor of two (from ∼10 to ∼5 mPa‧s). We also tested whether this altered condensate stability affects the partitioning of client molecules RP3 ((RRASL)_3_), an arginine-rich peptide that interacts with the condensates via electrostatic force^31^. The client RP3 exhibited a significant drop in partition coefficient from 30 to 8 after the addition of 0.9 M glycine (**Figure 1h**).

### Glycine dissolves synthetic condensates driven by electrostatic interactions and promotes those driven by cation-π and π-π interactions

To deconvolute the interplay of glycine with the complex multimodal interactions^32^ that drive the formation of NPM1-RNA condensates, four model synthetic condensates with simplified driving forces were employed. As shown in **Figure 2a**, the lysine(K)-rich elastin-like protein GFP-GFPGAGP[GVGVP(GKGVP)_9_]_8_GWPH_6_ (K72 in short), which contains 72 repeats of the pentapeptide GKGVP (an elastin-like sequence)^33^ fused to an N-terminal green fluorescent protein (GFP) for visualization purposes, can form condensates at low concentrations (10 µM) with ATP by electrostatic interactions under physiological pH (25 mM HEPES, pH 7.4)^20^. We measured the K72 concentration in the dilute phase by the fluorescence emission from the conjugated GFP and we found that with higher glycine concentration in solution, K72 concentrations in the dilute phase increased (**Figure 2a**). This agreed with a lower K72 partitioning in the condensate phase (**Figure S4**), which both indicate a gradual condensate dissolution process with the addition of glycine. By measuring protein concentration changes in the dilute phase, we found that the effect of glycine on the phase behavior of K72-ATP is qualitatively similar to that of NPM1-RNA (**Figure S5**) although they are dominated by different condensate-driving forces.

**Figure 2:**
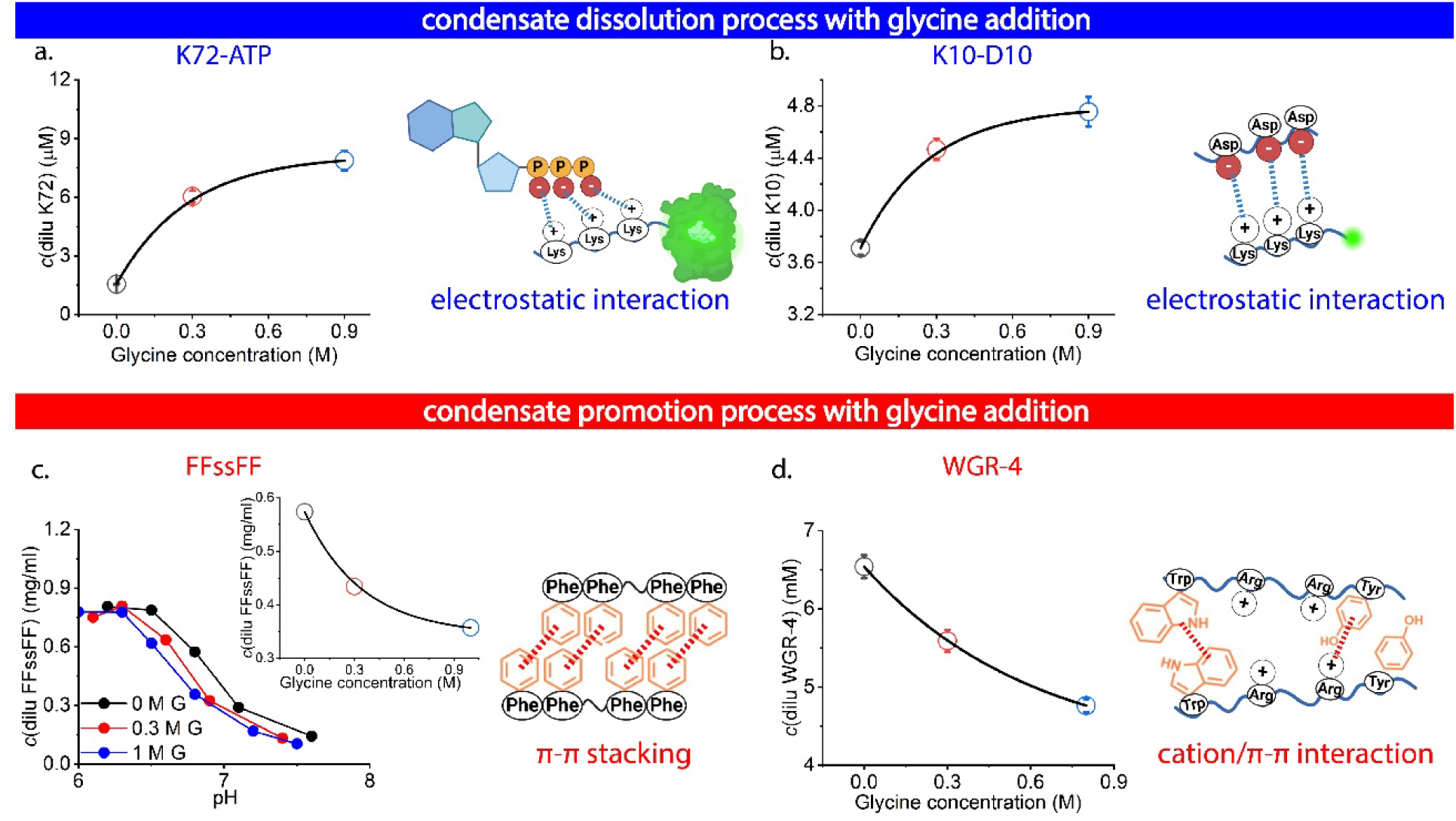
Model synthetic condensates to deconvolute the complex interaction in NPM1-RNA condensates. The peptide/protein concentration in the dilute phase after the addition of glycine (0, 0.3, and 0.9 M) as well as the expected intermolecular interactions underlying the condensate formation of a. K72-ATP system; b. K10-D10 system; c. FFssFF system (the inlet shows the FFssFF concentration in the dilute phase at pH 6.8) and d. WGR-4 system.

A second synthetic model condensate that is driven by electrostatic interaction consists of two short peptides, oligo-lysine (K_10_) and oligo-aspartic acid (D_10_)^21^. We measured the K_10_ concentration in the dilute phase by the fluorescence emission from the fluorescein-labeled K_10_ and found again that with higher glycine concentration in solution, K_10_ concentration in the dilute phase increased, which also indicates a similar condensate dissolution process (**Figure 2b**). This also agreed with a lower K_10_ partitioning in the condensate phase (**Figure S6**).

We then switched to model condensates formed by π-π stacking interactions and cation-π interactions, which are also among the major driving forces for condensate formation^6,34^. We chose a minimal sticker and spacer architecture model in the form of FFssFF, which contains two diphenylalanine stickers^35^. FFssFF molecules are soluble in acidic pH due to their net positive charge and start to form condensates due to π-π stacking when the pH is increased above approximately 6.5. We found that with increasing glycine concentration, the concentration of FFssFF in the dilute phase decreased, and condensates started to form at a lower pH (**Figure 2c**). Accordingly, the turbidity of the whole phase-separating solution increased at a lower pH (**Figure S7**), which indicates a promotion of condensate formation with the presence of glycine. Another minimalistic homotypic peptide condensate based on cation-π and π-π interactions was employed. The decapeptide WGR-4 (W(GR)_3_GWY) was reported by Lampel and co-workers^23^ to form condensates at neutral pH due to cation-π attractions between arginine residues (R) and aromatic residues (W and Y) as well as π-π stacking among aromatic residues (W and Y). Similar to the FFssFF system, we also found that with increasing glycine concentration, the WGR-4 concentration in the dilute phase decreased (**Figure 2d**) and the turbidity of the solution increased (**Figure S8**), which both indicates a promotion of condensate formation with the addition of glycine, similar to the FFssFF system.

### Backbone and aryl binding of amino acids underlie the modulation of condensate stability

The curves of increasing dilute phase NPM1 concentration with increasing glycine concentration are reminiscent of a binding isotherm. We used a Langmuir-type binding model^36^ to fit the NPM1 protein concentration change in the dilute phase Δ*c*(dilu NPM1) as a function of glycine concentration and found an apparent dissociation constant (*K*_d_) of glycine of ∼0.9 M (**Figure 3a**), corresponding to weak affinity binding. This weak apparent binding between glycine and the condensates agrees with the recently proposed theory that AAs can weakly bind to proteins and modulate protein-protein interactions by effectively screening a fraction of their net interaction potential^15^. We also measured the binding affinity of four different AAs (glycine/G, proline/P, serine/S, and alanine/A) to K72-ATP condensates (K72 concentration change in the dilute phase as a function of AA concentration in **Figure S9**). All the fitted values of *K*_d_ were found to be ∼1 M (**Figure 3b**).

**Figure 3:**
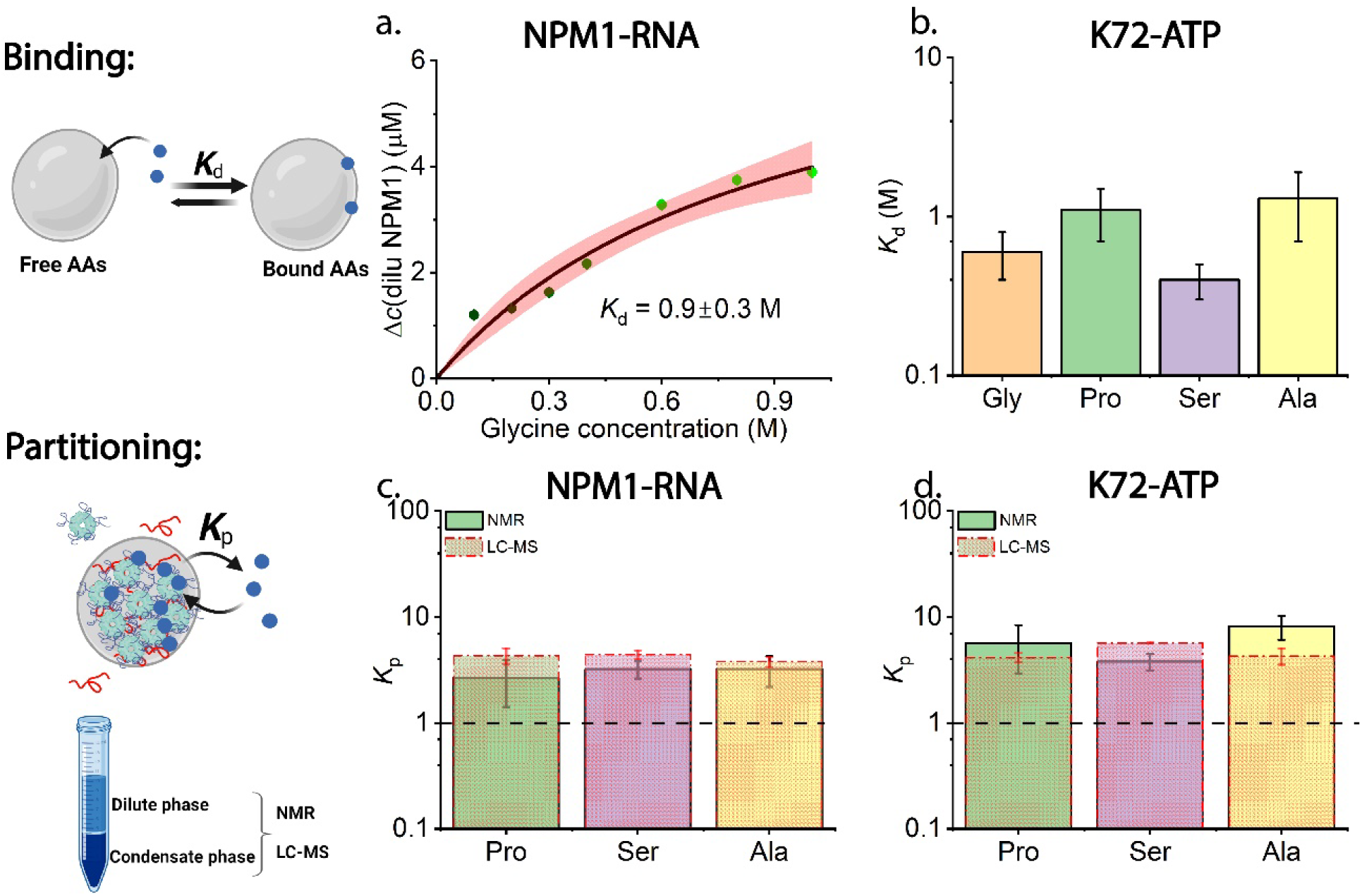
Binding and partitioning of AAs in NPM1-RNA and K72-ATP condensates. a. The NPM1 protein concentration in the dilute phase after adding glycine at varying concentrations and the fitting curve (black) with 95% confidence band (red) using the simple Langmuir-type binding model; b. The fitted binding affinities (*K*_d_) for four different AAs (glycine, proline, serine, and alanine) on K72-ATP condensates; The measured partition coefficients (*K*_p_) by both NMR and LC-MS for three representative AAs (proline, serine, and alanine) in the condensate phase of c. NPM1-RNA systems and d. K72-ATP systems.

Weak binding of AAs to NPM1, K72, and possibly other phase-separating proteins suggests that AAs should be accumulated in the condensates. We therefore employed LC-MS and ^1^H NMR to measure the partition coefficients (*K*_p_) of AAs.^37–39^ We found that *K*_p_ for all the tested AAs are ∼4 in NPM1-RNA condensates and ∼6 in K72-ATP condensates (NMR spectrum with peak assignments in **Figure S10** and LC-MS raw data in **Figure S11**), in reasonable agreement with an apparent dissociation constant of 0.9 M and an estimated local protein concentration of 1 mM for K72^20^.

The Langmuir-type binding model suggests that AAs bind to specific sites along the protein. To elucidate the binding positions of AAs on BCs, we employed NMR spectroscopy under conditions where phase separation does not occur. Following the assignments of proton peaks by the Total Correlation Spectroscopy (TOCSY) (**Figure S12**), we ran ^1^H NMR experiments by the titration of deuterium-labelled glycine (G-d_5_) into solutions of K72-GFP. We observed significant changes in backbone amide chemical shifts of glycine/G and valine/V residues in K72 while the chemical shifts for the other proton peaks hardly changed, which indicates the binding of G to backbone amide groups (**Figure 4a**). The chemical shift perturbation (CSP) at different concentrations of G-d_5_ was also fitted by the Langmuir-type binding model^36^ and the dissociation constant (*K*_d_) of G-d_5_ and amide groups in G and V residues was estimated to be 1.5 ± 0.2 and 1.7 ± 0.3 M respectively (**Figure 4d**). The overall *K*_d_ of G-d_5_ to K72 can be estimated to be ∼0.8 M by a first-order approximation, which agrees well with the binding affinity obtained from Δ*c*(K72) in the dilute phase (0.6 ± 0.2 M) (**Figure 3b**).

**Figure 4:**
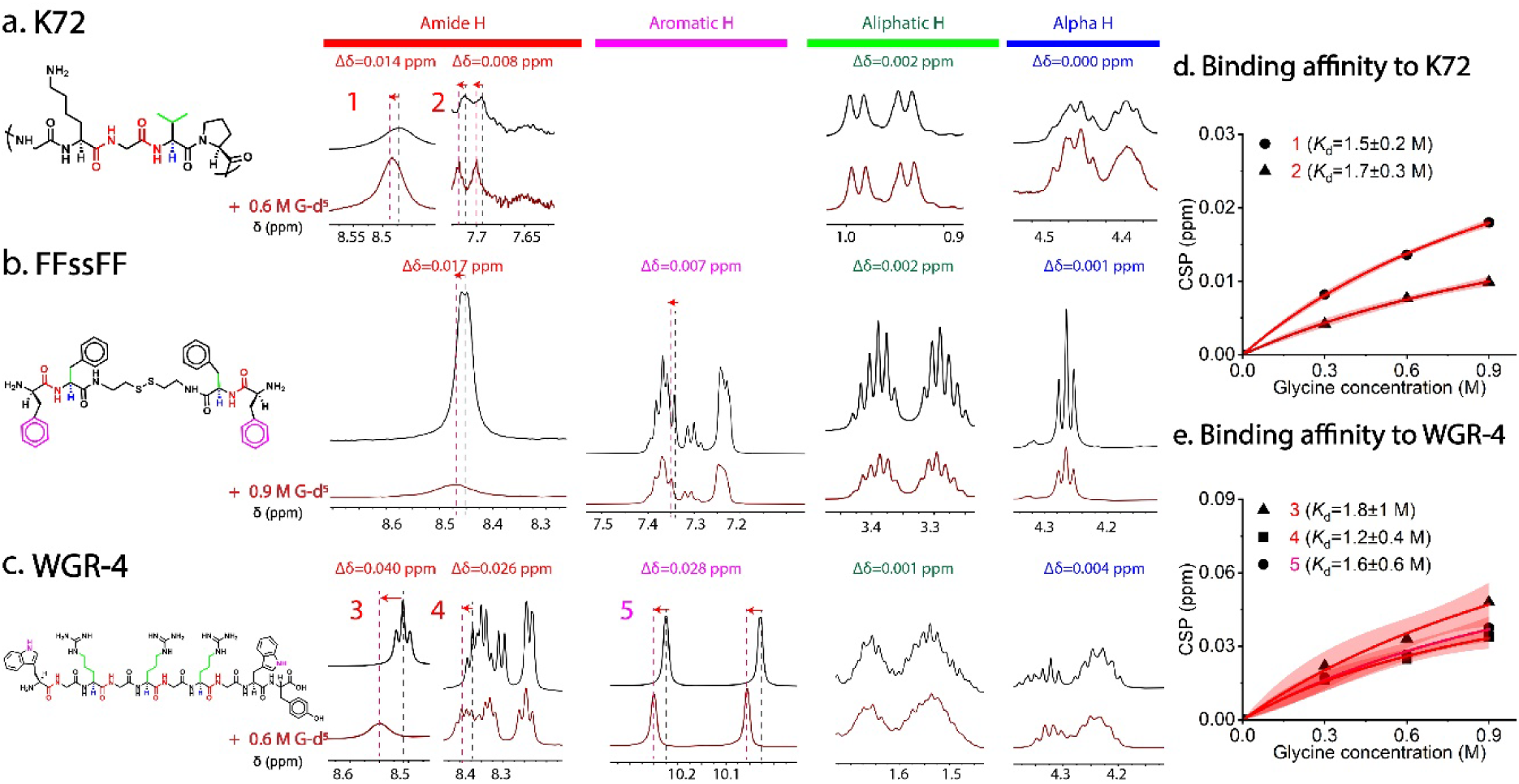
Binding of glycine (G-d_5_) to different proteins/peptides from ^1^H NMR spectroscopy. The chemical structures, the proton peaks that experience significant chemical shifts (in red and pink) and lack significant chemical shift perturbations (in green and blue) with (red spectrum) and without (black spectrum) G-d_5_ for a. K72; b. FFssFF and c. WGR-4; The chemical shift perturbation (CSP) of d. K72 and e. WGR-4 with the titration of G-d_5_ at 4 different concentrations to estimate the binding affinity (*K*_d_) of G to K72 and WGR-4. The fitting curves are in black with 95% confidence band (red) using the simple Langmuir-type binding model.

Interestingly, the ^1^H NMR experiments of FFssFF and WGR-4 after the addition of G-d_5_ also show significant changes in backbone amide group chemical shifts. Moreover, we also observed significant changes in side-chain aromatic group chemical shifts (**Figure 4b** and **4c**). The chemical shifts for the other groups remained unchanged (TOCSY in **Figure S13** for the amide and aromatic group assignments). The binding affinity of glycine to aromatic groups is also similar to that of amide groups, both characterized by an apparent *K*_d_ of ∼1.5 M (**Figure 4e**). Overall, the chemical shift data of these peptides show a consistent perturbation of backbone amide groups by glycine, which impacts inter-molecular amide-amide hydrogen bonds. The additional perturbation on side chain aromatic groups is observed for peptides rich in aromatic AAs, which has additional effects on inter-molecular π-π stacking and cation-π interactions.

### Condensate modulation extends to most proteogenic amino acids and is transferable to short peptides

To investigate whether the modulation effect of glycine is a general property of AAs, we screened all the neutral proteogenic AAs with a solubility > 100 mM as well as charged AAs for their effect on NPM1-RNA, K72-ATP, and WGR-4 condensate systems using a microplate reader assay. As shown in **Figure 5a**, all the AAs tested show a similar condensate dissolution effect to glycine (G) on NPM1-RNA, as indicated by higher protein concentrations in the dilute phase after the addition of AAs. The only exception is glutamic acid (E). We found slightly lower (∼2%) protein concentrations in the dilute phase after the addition of 20 mM sodium glutamate compared to the control set, indicating enhanced condensate formation. Similarly, except for glutamate (E), all the neutral AAs tested show a condensate dissolution effect on K72-ATP system. Interestingly, positively-charged AAs (H and K) show a more significant dissolution effect on K72-ATP condensates, which may be because those AAs compete with K72 for binding to negatively-charged ATP, weakening the electrostatic attractions governing the condensate formation. On the other hand, negatively charged AA glutamic acid (E) shows a significant promotion of K72-ATP condensate formation. Similar condensate promotion effects were also observed for all the tested proteinogenic AAs on WGR-4 systems, except for one outlier proline (**Figure S14**).

**Figure 5:**
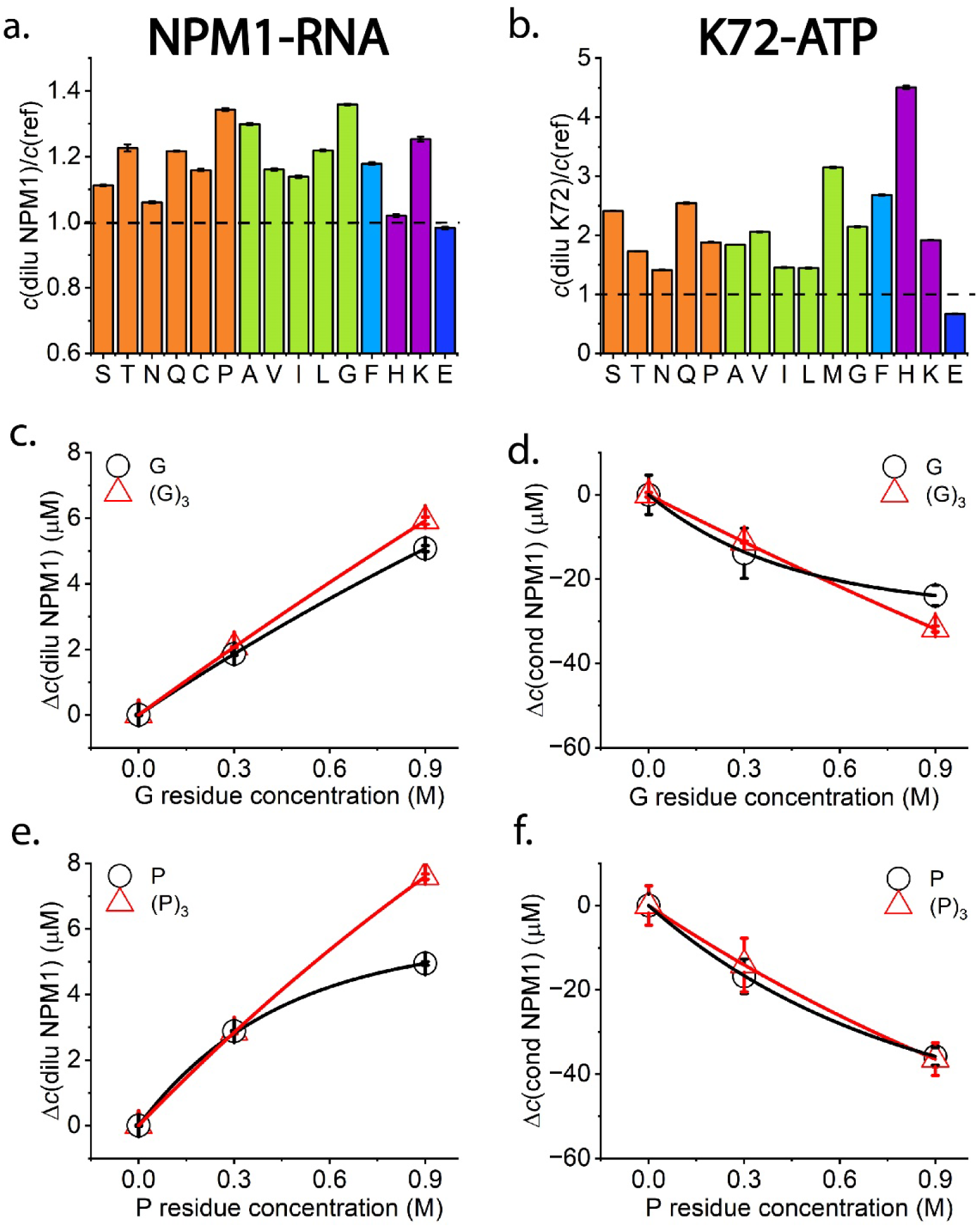
General modulation effects of AAs and the transferability to short peptides. a. NPM1 concentration in the dilute phase after the addition of different AAs (200 mM of S, T, Q, C, G, P, A, V; 100 mM of H, N, I, L, F; 20 mM K, E) divided by reference NPM1 concentration in the dilute phase in the absence of any AAs; b. K72 concentration in the dilute phase after the addition of different AAs (200 mM of S, T, Q, G, P, A, V; 100 mM of M, H, N, I, L, F; 20 mM K, E) divided by reference K72 concentration in the dilute phase in the absence of any AA; AAs are groups in colours by side chain properties. Orange, green blue and purple bars represent polar uncharged, nonpolar aliphatic, nonpolar aromatic and slightly positive charged side groups respectively; c-d. NPM1 concentration changes in the dilute and condensate phase after the addition of G and (G)_3_ at the same ionic strength; e-f. NPM1 concentration changes in the dilute and condensate phases after the addition of P and (P)_3_ at the same ionic strength.

Furthermore, two short peptides, a glycine trimer (G)_3_ and proline trimer (P)_3,_ were employed to investigate if they could also modulate condensate formation. As shown in **Figure 5c-f**, very similar modulation effects on the NPM1-RNA phase separation were observed for these two short homo-peptides at the same AA residue concentrations as free AAs, which indicates that the modulation effect is transferable to peptides in an additive manner^14^. Moreover, the linear transferability in the modulation effect is still true for proline octamer (P)_8_ (**Figure S15a and S15b**). However, aggregation started to appear (**Figure S15c and S15d**), which may be due to the molecular rigidity of a relatively long peptide chain compared to free AAs.

## Discussion

Based on the experimental data presented in this study, we propose a multiscale mechanism behind the macroscopic modulation effect of AAs on BCs, **Figure 6**. On a molecular scale, AAs bind to backbone amide groups of phase-separating proteins, which weaken the intermolecular amide-amide hydrogen bonds. This explains that AAs can dissolve NPM1-RNA, K72-ATP, and K10-D10 condensates. We also find that most AAs have similar modulation effects (**Figure 5**). This suggests that the binding is mediated primarily through the free amine or carboxylic acid group in the AAs’ main chain.

**Figure 6:**
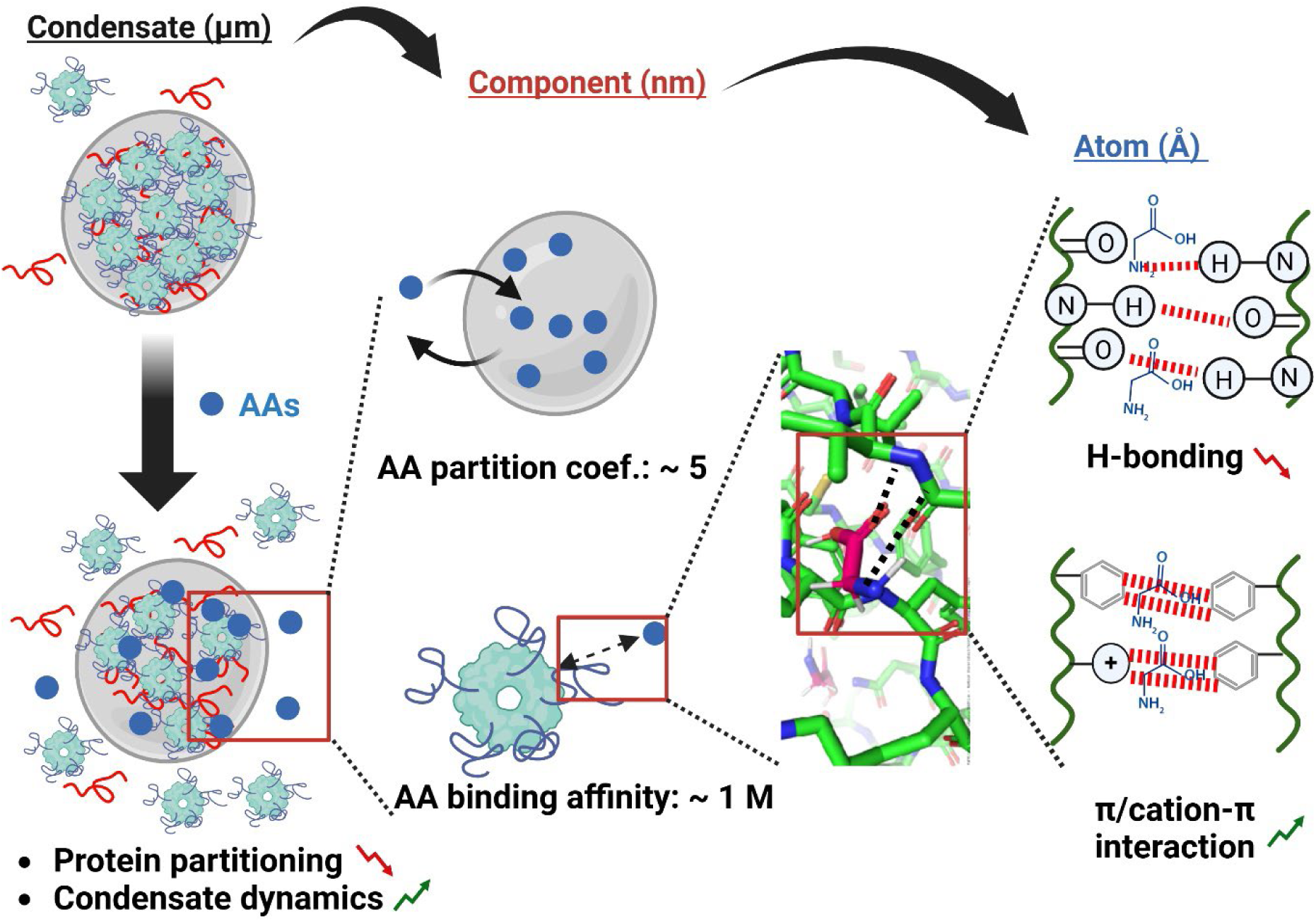
The proposed multiscale mechanism for the modulation effect of AAs on BCs, ranging from µm-scale condensate formation and dynamics, nm-scale component interaction and partitioning to Å-scale atomic interaction. Created by Biorender.

To shed more light on the binding mechanism, we investigated two glycine derivatives: betaine, which contains a tertiary amine instead of a primary amine, and taurine, which contains a sulfonic acid instead of a carboxylic acid group. We found that adding betaine did not affect NPM1-RNA condensate formation while adding taurine similarly dissolved condensates as glycine (**Figure S16**). This indicates that the primary amine group of AAs most likely binds to the proteins in BC components through hydrogen bonding.

In addition, we employed synthetic condensates formed by the electrostatic interaction between poly(diallyldimethyl-ammonium chloride) (PDDA) and poly(acrylic acid) (PAA), that do not contain amide groups in their backbones. We found that the addition of glycine did not affect the phase separation of this system (**Figure S17**). This experiment further supports our hypothesis that AAs bind to amide groups in the backbone of disordered proteins and modulate the protein backbone-backbone interactions, which contribute to condensate stability. AAs do not suppress phase separation when there are no amide groups in their backbone to bind to, such as in the case of PDDA-PAA condensates. In addition, the binding of AAs to side-chain aromatic groups can increase the polarity around aromatic rings, which may enhance cation/π-π interaction^25,26,40^. This explains that AAs can promote the formation of aromatic-rich FFssFF and WGR-4 condensates.

According to our proposed molecular mechanism, no specific AA side chains are needed for the observed interaction with BC components. This is supported by our findings that most AAs have similar modulation effects (**Figure 5**). More importantly, the observed transferability of the modulation to short peptides indicates that short peptides can bind reasonably strongly to disordered protein backbones, and opens the way for the rational design of short peptides to modulate BCs by strategically engineering side-chain interactions at specific binding sites. We can harness this biocompatible and versatile peptide platform to specifically target the properties and functions of physiologically important biomolecular condensates in the future^41–43^. These strategies may help to control the biological processes related to BCs, and potentially suppress undesired aging of condensates, which has been linked to a variety of neurodegenerative diseases^8,44^.

## Methods

### Materials

All chemicals and reagents were used as received from commercial suppliers. The following chemicals were purchased from Sigma Aldrich: adenosine triphosphate (ATP), sodium chloride, Tris base, PEG(10k Da), and all the 18 amino acids that are used in the study, including G, S, T, Q, C, P, A, V, M, H, N, I, L, F, R, K, D and E and glycine-d^5^ (175838). poly(acrylic acid) sodium salt (PAA, 15 kDa, 35 wt % solution in H2O) and poly(diallyldimethylammonium chloride) (PDDA, 200–350 kDa, 20 wt % solution in H2O).HEPES-free acid was purchased from FluoroChem. PLL-*g*[3.5]-PEG was purchased from SuSoS. Poly-L-lysine hydrobromide (MW = 2100 Da, 10-mer) and poly-L-aspartic acid sodium salt (MW = 1400 Da, 10-mer) were purchased from Alamanda Polymers. WGR-4, (G)3, (P)3, and (P)8 were all purchased from Genscript Biotech (The Netherlands, The Hague). 5,6-FAM-RP3 was purchased from CASLO, Kongens Lyngby, Denmark. GFP-labelled K72 and NPM1 were expressed and purified as previously described^19,20^. FFssFF was synthesized as previously described^35^. *E. coli* rRNA was purified as previously described^19^.

### Condensate formation

NPM1-RNA condensates were prepared in Tris buffer (final concentration 10 mM, pH 7.5) with 150 mM NaCl, by adding PEG 10k Da (final concentration 2.3 wt%) and RNA/RNA-A647 (final concentration 100 ng/μL, 1:19 molar ratio labelled) to Tris buffer followed by NPM1/NPM1-A488 (final concentration 20 µM, 1:19 molar ratio labelled). K72-ATP condensates were prepared in HEPES buffer (final concentration 25 mM, pH 7.4), by adding GFP-labelled K72 (final concentration 10 µM) to HEPES buffer followed by ATP (final concentration 4 mM). K10-D10 condensates were prepared in HEPES buffer (final concentration 50 mM, pH 7.4), by adding D10 (final concentration 5 mM) to HEPES buffer followed by fluorescein labelled K10 (final concentration 5 mM). 10 mg/ml FFssFF stock solution in water was diluted 10 times with pH buffers of the pH values from 5 to 8. 9 mM WGR-4 stock solution in water was added into the Tris buffer (final concentration 20 mM, pH 7.5) with 800 mM NaCl. AAs of specified final concentrations were prepared in the buffer solution by adding specific volumes of AA stock solutions before the proteins/peptides were added.

### Confocal fluorescence microscopy

All samples were imaged on Ibidi 18-well chambered slides (#1.5) that were cleaned with a plasma cleaner, incubated for 24 h with 0.1 mg mL−1 PLL-g[3.5]-PEG (SuSoS, Dübendorf, Switzerland) dissolved in Milli-Q water, and washed and dried with Milli-Q water and pressurized air, respectively. Before image acquisition, the condensate dispersion was transferred to the channel and incubated for over 0.5 h to allow condensates to coalesce and settle on the glass surface. Confocal fluorescence images were acquired on a Leica Sp8x confocal inverted microscope (Leica Microsystems, Germany) equipped with a DMi8 CS motorized stage, a pulsed white light laser, and 2 × HyD SP GaAsP and 2× PMT detectors. Images were recorded using the LAS X v.3.5 acquisition software, using an HC PL APO 100×/1.40 oil immersion objective.

The calculation of partition coefficients was automated with a MATLAB script for at least 3 fluorescence images by the following equation: *K*p = (*I*condensate – *I*background)/(*I*dilute – *I*background), where *I*condensate denotes the average intensity of condensates in one image, and *I*dilute denotes the average intensity of the area without condensates, *I*background denotes the background intensity by measuring only the buffer at the same settings as for the fluorescent images, which is normally zero.

### Quantification of the protein concentrations in the dilute and condensate phases

A typical sample of 38 μL was prepared in 10 mM Tris (pH 7.5) and 150 mM NaCl with 2.3 wt% of PEG, 20 µM NPM1/NPM1-A488 (1:19 molar ratio labelled), and 100 ng/μL RNA (unlabelled) as described above. After the incubation for 20 min at room temperature, the condensate phase was separated from the dilute phase by centrifugation at 20,000 g for 20 min at room temperature. The supernatant of 20 μL was then transferred to a 384-well plate (Nunc, flat bottom), and the fluorescence intensity was measured on a plate reader (Tecan Spark M10) at 485/535 nm for NPM1-A488 and 620/680 nm for RNA-A647. Concentrations of the dilute phase were calculated based on calibration curves (**Figure S18**). The NPM1 concentration in the condensate phase can be then calculated by the following equation: *c*(NPM1 in condensate phase) = c(NPM1 in dilute phase) × *K*p.

### Fluorescence recovery after photobleaching

Time-lapse videos were recorded at room temperature on a CSU X-1 Yokogawa spinning disk confocal unit connected to an Olympus IX81 inverted microscope, using an ×100 piezo-driven oil immersion objective (NA 1.3) and 488 and 640 nm laser beams. Emission was measured with a 200-ms exposure time using an Andor iXon3 EM-CCD camera. The acquired images have a pixel size of 141 nm and a field of 512 x 512 μm^2^. For the laser bleaching, a small region of interest was selected in the middle of a condensate droplet. The 488 or 640 nm laser line was set to 100% laser power using 20 pulses of 200 μs. The recovery was then imaged at reduced laser intensity with a time interval of 300 ms for 200 times. The exponential decay equation *I*normalized = *A*(1-e^-*bt*^)+*C* was used to fit the parameters *A*, *b*, and *C,* and the recovery half-life was determined by the equation: *t*1/2 = *ln*(2)/*b* according to a 2D-diffusion model with a fixed boundary^45^.

### Diffusion coefficients measured by Raster image correlation spectroscopy (RICS)

The Raster Image Correlation Spectroscopy (RICS) was performed on a Leica SP8 confocal microscope equipped with a single-photon detector. Calibration of the focal volume waist *ω*0 was performed using the known diffusion coefficient of Alexa 488 of 435 μm^2^/s (T = 22.5 ± 0.5 °C) in water, and *ω*z was set to 3 times the value of *ω*_0_ ^46^. All measurements were captured at a resolution of 256 × 256 pixels with a 20 nm pixel size using a 63x oil objective. Condensates were measured at 10 Hz line speed with 15 frames acquired per data point. Analysis of autocorrelation curves was done using PAM^47^.

### Langmuir-type binding model fitting of dilute phase NPM1 concentration change and the chemical shift perturbation with different glycine concentrations

The dilute phase NPM1 concentration change and the chemical shift perturbation (both denoted as *Δ*) with different AA concentrations (*c*) was fitted by a simple binding model^15,36^ under the assumption of excess AA: 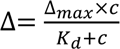.

### Sample preparation for the partitioning coefficients measurements of AAs

In a PCV cell counting tubes (capillary graduations only, no cap, Sigma-Adrich), NPM1-RNA condensates were prepared in Tris buffer (final concentration 10 mM, pH 7.5) with 150 mM NaCl, by adding PEG 10k Da (final concentration 2.3 wt%) and RNA (final concentration 100 ng/μL) to Tris buffer followed by NPM1 (final concentration 20 µM) at a total sample volume of 400 μL. After the incubation at RT for 30 mins, the tube was centrifuged at 3200 g for 30 min at RT to spin down the condensate phase. supernatant (dilute phase) was carefully separated from the viscous condensate phase and stored in a separate tube. The condensate phase was redispersed with 1xPBS in an appropriate volume ratio. The dilute phase was diluted with 1xPBS in the same ratio. All the macromolecules were removed by using Amicon^®^ Ultracentrifuge Fiter units (10k Da cut-off) at 4000 g for 30 min. The filtered solution was ready for the partition coefficient measurements by NMR or LC-MS.

### Partitioning coefficient and chemical shift perturbation experiments by NMR

NMR samples were prepared by dissolving proteins or peptides (10 µM of K72 and 1 mM of WGR-4 proteins) in 500 µl of 1×PBS buffer (pH 7.2) with 10% D2O (containing 0.05 wt.% 3-(trimethylsilyl)propionic-2,2,3,3-d4 acid, sodium salt as the internal standard). Measurements were conducted on a Bruker Avance III 500 MHz NMR Spectrometer equipped with a Prodigy BB cryoprobe at 298.15 K. 1D-^1^H experiments were performed using the zgesgp water-suppression pulse sequence with 128 scans and a total relaxation and acquisition of 6.3 s. For the chemical shift perturbation experiments, 1D-^1^H experiments were performed using the zgesgp water-suppression pulse sequence with 128 scans and a total relaxation and acquisition of 6.3 s and 2D-^1^H,^1^H-TOCSY experiments were performed with 60 ms spin-lock, 64 scans per increment, 512 increments with a 6 kHz spectral window in dimension.

### Partitioning coefficient measurements of AAs by LC-MS

Proline, serine and alanine concentrations were determined using an Agilent 1290 Infinity II LC system coupled to an Agilent Accurate Mass 6546 Quadrupole - Time of Flight (Q-TOF) mass spectrometer, using a previously described method^48^. In brief, 2 µL samples were injected onto a Diamond Hydride Type C column (Cogent) and separated using a 0.4 mL/min gradient of water with 0.2% formic acid (A) in acetonitrile with 0.2% formic acid (B) as follows: 0–2 min: 85% B, 3–5 min: 80% B, 6–7 min: 75% B, 8-9 min: 70% B, 10–11 min: 50% B, 11–14: 20% B, 14–24: 5% B, followed by 10 min re-equilibration at 85% B. Detection was performed in the positive ionization mode and a mass range of m/z 50-1200. Analyte peaks were extracted using a 20 ppm window and integrated manually.

## Acknowledgements

The authors acknowledge dr. R.M. de Graaf for running the LC-MS. X.F.X acknowledges the Swiss National Science Foundation for financial support (P500PN_222304). E.S. acknowledges funding from a Vidi grant from the Netherlands Organization for Scientific Research (NWO).

## References

1. Spruijt, E. Open questions on liquid–liquid phase separation. Commun Chem 6, 1–5 (2023).

2. Yewdall, N. A., André, A. A. M., Lu, T. & Spruijt, E. Coacervates as models of membraneless organelles. Current Opinion in Colloid & Interface Science 52, 101416 (2021).

3. Astoricchio, E., Alfano, C., Rajendran, L., Temussi, P. A. & Pastore, A. The Wide World of Coacervates: From the Sea to Neurodegeneration. Trends in Biochemical Sciences 45, 706– 717 (2020).

4. Pederson, T. The nucleolus. Cold Spring Harb Perspect Biol 3, a000638 (2011).

5. Brangwynne, C. P. et al. Germline P Granules Are Liquid Droplets That Localize by Controlled Dissolution/Condensation. Science 324, 1729–1732 (2009).

6. Wang, J. et al. A Molecular Grammar Governing the Driving Forces for Phase Separation of Prion-like RNA Binding Proteins. Cell 174, 688–699.e16 (2018).

7. Li, P. et al. Phase transitions in the assembly of multivalent signalling proteins. Nature 483, 336–340 (2012).

8. Alberti, S. & Hyman, A. A. Biomolecular condensates at the nexus of cellular stress, protein aggregation disease and ageing. Nat Rev Mol Cell Biol 22, 196–213 (2021).

9. Banani, S. F., Lee, H. O., Hyman, A. A. & Rosen, M. K. Biomolecular condensates: organizers of cellular biochemistry. Nat Rev Mol Cell Biol 18, 285–298 (2017).

10. Mitrea, D. M., Mittasch, M., Gomes, B. F., Klein, I. A. & Murcko, M. A. Modulating biomolecular condensates: a novel approach to drug discovery. Nat Rev Drug Discov 21, 841–862 (2022).

11. André, A. A. M. & Spruijt, E. Liquid–Liquid Phase Separation in Crowded Environments. IJMS 21, 5908 (2020).

12. Patel, A. et al. ATP as a biological hydrotrope. Science 356, 753–756 (2017).

13. Rollin, R., Joanny, J.-F. & Sens, P. Physical basis of the cell size scaling laws. eLife 12, e82490 (2023).

14. Xu, X. & Stellacci, F. Amino Acids and Their Biological Derivatives Modulate Protein– Protein Interactions in an Additive Way. J. Phys. Chem. Lett. 15, 7154–7160 (2024).

15. Mao, T., et al. Amino Acids Stabilizing Effect on Protein and Colloidal Dispersions. Preprint at 10.48550/arXiv.2404.11574 (2024).

16. Xu, X. et al. Amino acids modulate liquid–liquid phase separation in vitro and in vivo by regulating protein–protein interactions. Proceedings of the National Academy of Sciences 121, e2407633121 (2024).

17. Paccione, G. et al. Lipid Surfaces and Glutamate Anions Enhance Formation of Dynamic Biomolecular Condensates Containing Bacterial Cell Division Protein FtsZ and Its DNA-Bound Regulator SlmA. Biochemistry 61, 2482–2489 (2022).

18. Kozlov, A. G. et al. How Glutamate Promotes Liquid-liquid Phase Separation and DNA Binding Cooperativity of E. coli SSB Protein. Journal of Molecular Biology 434, 167562 (2022).

19. Yewdall, N. A. et al. ATP:Mg2+ shapes material properties of protein-RNA condensates and their partitioning of clients. Biophysical Journal 121, 3962–3974 (2022).

20. Nakashima, K. K., van Haren, M. H. I., André, A. A. M., Robu, I. & Spruijt, E. Active coacervate droplets are protocells that grow and resist Ostwald ripening. Nat Commun 12, 3819 (2021).

21. van Haren, M. H. I., Visser, B. S. & Spruijt, E. Probing the surface charge of condensates using microelectrophoresis. Nat Commun 15, 3564 (2024).

22. Lipiński, W. P., et al. Biomolecular condensates can both accelerate and suppress aggregation of α-synuclein. Science Advances 8, eabq6495 (2022).

23. Baruch Leshem, A., et al. Biomolecular condensates formed by designer minimalistic peptides. Nat Commun 14, 421 (2023).

24. Tsuzuki, S. Interactions with Aromatic Rings. in Intermolecular Forces and Clusters I (ed. Wales, D. J.) 149–193 (Springer, Berlin, Heidelberg, 2005). doi:10.1007/b135618.

25. Hallett, J. E., Agg, K. J. & Perkin, S. Zwitterions fine-tune interactions in electrolyte solutions. Proceedings of the National Academy of Sciences 120, e2215585120 (2023).

26. Mei, W. et al. Zwitterions Raise the Dielectric Constant of Soft Materials. Phys. Rev. Lett. 127, 228001 (2021).

27. Govrin, R., Tcherner, S., Obstbaum, T. & Sivan, U. Zwitterionic Osmolytes Resurrect Electrostatic Interactions Screened by Salt. J. Am. Chem. Soc. 140, 14206–14210 (2018).

28. Mitrea, D. M. et al. Self-interaction of NPM1 modulates multiple mechanisms of liquid– liquid phase separation. Nat Commun 9, 842 (2018).

29. André, A. A. M., Yewdall, N. A. & Spruijt, E. Crowding-induced phase separation and gelling by co-condensation of PEG in NPM1-rRNA condensates. Biophysical Journal 122, 397–407 (2023).

30. Qian, D. et al. Tie-Line Analysis Reveals Interactions Driving Heteromolecular Condensate Formation. Phys. Rev. X 12, 041038 (2022).

31. Musinova, Y. R., Kananykhina, E. Y., Potashnikova, D. M., Lisitsyna, O. M. & Sheval, E. V. A charge-dependent mechanism is responsible for the dynamic accumulation of proteins inside nucleoli. Biochimica et Biophysica Acta (BBA) - Molecular Cell Research 1853, 101–110 (2015).

32. Mitrea, D. M. et al. Nucleophosmin integrates within the nucleolus via multi-modal interactions with proteins displaying R-rich linear motifs and rRNA. eLife 5, e13571 (2016).

33. Yang, H. et al. Tuning Ice Nucleation with Supercharged Polypeptides. Advanced Materials 28, 5008–5012 (2016).

34. Brangwynne, C. P., Tompa, P. & Pappu, R. V. Polymer physics of intracellular phase transitions. Nature Phys 11, 899–904 (2015).

35. Abbas, M., Lipiński, W. P., Nakashima, K. K., Huck, W. T. S. & Spruijt, E. A short peptide synthon for liquid–liquid phase separation. Nat. Chem. (2021) doi:10.1038/s41557-021-00788-x.

36. Milles, S. et al. Plasticity of an ultrafast interaction between nucleoporins and nuclear transport receptors. Cell 163, 734–745 (2015).

37. Soininen, P. et al. High-throughput serum NMR metabonomics for cost-effective holistic studies on systemic metabolism. Analyst 134, 1781–1785 (2009).

38. Ambadi Thody, S., et al. Small-molecule properties define partitioning into biomolecular condensates. Nat. Chem. 1–9 (2024) doi:10.1038/s41557-024-01630-w.

39. Wang, J., Abbas, M., Wang, J. & Spruijt, E. Selective amide bond formation in redox-active coacervate protocells. Nat Commun 14, 8492 (2023).

40. Azizi, A. & Ebrahimi, A. Theoretical investigation of the π+-π+ stacking interactions in substituted pyridinium ion. Journal of Molecular Graphics and Modelling 77, 225–231 (2017).

41. Alberti, S., Gladfelter, A. & Mittag, T. Considerations and Challenges in Studying Liquid-Liquid Phase Separation and Biomolecular Condensates. Cell 176, 419–434 (2019).

42. Luo, J. & Abrahams, J. P. Cyclic Peptides as Inhibitors of Amyloid Fibrillation. Chemistry – A European Journal 20, 2410–2419 (2014).

43. Merz, M. L. et al. De novo development of small cyclic peptides that are orally bioavailable. Nat Chem Biol 1–10 (2023) doi:10.1038/s41589-023-01496-y.

44. Patel, A. et al. A Liquid-to-Solid Phase Transition of the ALS Protein FUS Accelerated by Disease Mutation. Cell 162, 1066–1077 (2015).

45. Taylor, N. O., Wei, M.-T., Stone, H. A. & Brangwynne, C. P. Quantifying Dynamics in Phase-Separated Condensates Using Fluorescence Recovery after Photobleaching. Biophysical Journal 117, 1285–1300 (2019).

46. Petrášek, Z. & Schwille, P. Precise Measurement of Diffusion Coefficients using Scanning Fluorescence Correlation Spectroscopy. Biophysical Journal 94, 1437–1448 (2008).

47. Schrimpf, W., Barth, A., Hendrix, J. & Lamb, D. C. PAM: A Framework for Integrated Analysis of Imaging, Single-Molecule, and Ensemble Fluorescence Data. Biophysical Journal 114, 1518–1528 (2018).

48. Jansen, R. S. et al. Aspartate aminotransferase Rv3722c governs aspartate-dependent nitrogen metabolism in Mycobacterium tuberculosis. Nat Commun 11, 1960 (2020).

