## Supplemental information for "Amino acids bind to phase-separating proteins and modulate biomolecular condensate stability and dynamics"

To further quantify the effect of glycine on intermolecular interactions underlying phase separation, we calculated the tie-line gradient *k,* according to Qian et al[30], which can be expressed by effective interaction difference for the condensate formation ($\chi^{\Delta}$): $k \approx -\frac{1}{(1+2\chi^{\Delta})N_1\phi_1}$ where $\phi_1$ and $N_1$ denote the volume fraction and length of the component 1 (NPM1), respectively, and $\chi^{\Delta} = \frac{z}{2k_BT}(\mu_{12} + \mu_{00} - \mu_{01} - \mu_{02})$, with $\mu$ the contact energy between the solvent (0), solute 1 (NPM1) and 2 (RNA), respectively, $z$ a coordination constant, $k_B$ Boltzmann's constant and $T$ the absolute temperature[30]. As shown in **Figure 1e**, the tie-line gradient *k* increases with increasing glycine concentration, indicating a weaker associative interaction driving the condensate formation.

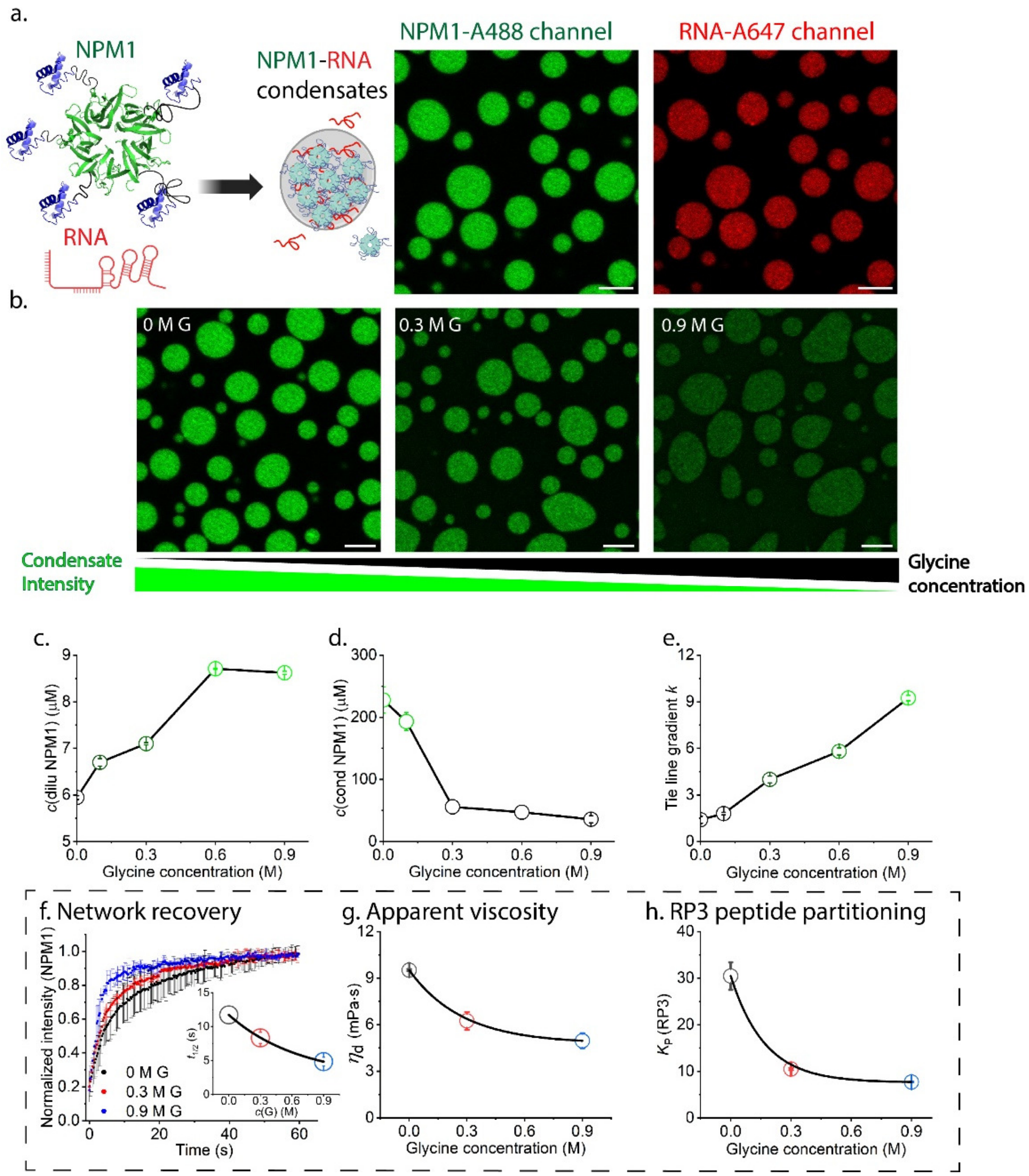


**Figure 1**: The phase behaviour and material properties of NPM1-RNA condensate after the addition of glycine. a. Schematic illustration of NPM1 protein structures (oligomerization domain (green, PDB: 4N8M) connected via disordered regions (grey) to the C-terminal nucleic acid binding domain (blue, PDB: 2VXD) and their formation of condensates with RNA. Fluorescence confocal microscopy images of NPM1-RNA condensates in NPM1-Alexa488 channel and RNA-Alexa647 channel; Scale bar = 5 µm. b. Confocal fluorescence microscopy images of NPM1-RNA condensates in NPM1-Alexa488 channel after the addition of 0, 0.3, and 0.9 M glycine; c and d. NPM1 concentrations in the dilute and condensate phases after the addition of glycine (0, 0.1, 0.3, 0.6, and 0.9 M) (calculation details in **Methods**); e. Calculated tie line gradient *k* after the addition of glycine (0, 0.1, 0.3, 0.6, and 0.9 M); f. Average FRAP recovery curves of NPM1 and the calculated recovery half-life ($t_{1/2}$) after the addition of glycine (0, 0.3, and 0.9 M); g. Apparent viscosity ($\eta_d$) of fluorescein (Alexa Fluor 488) in NPM1-RNA condensates after the addition of glycine (0, 0.3, and 0.9 M); h. Partitioning coefficients ($K_p$) of RP3 in NPM1-RNA condensates after the addition of glycine (0, 0.3, and 0.9 M).

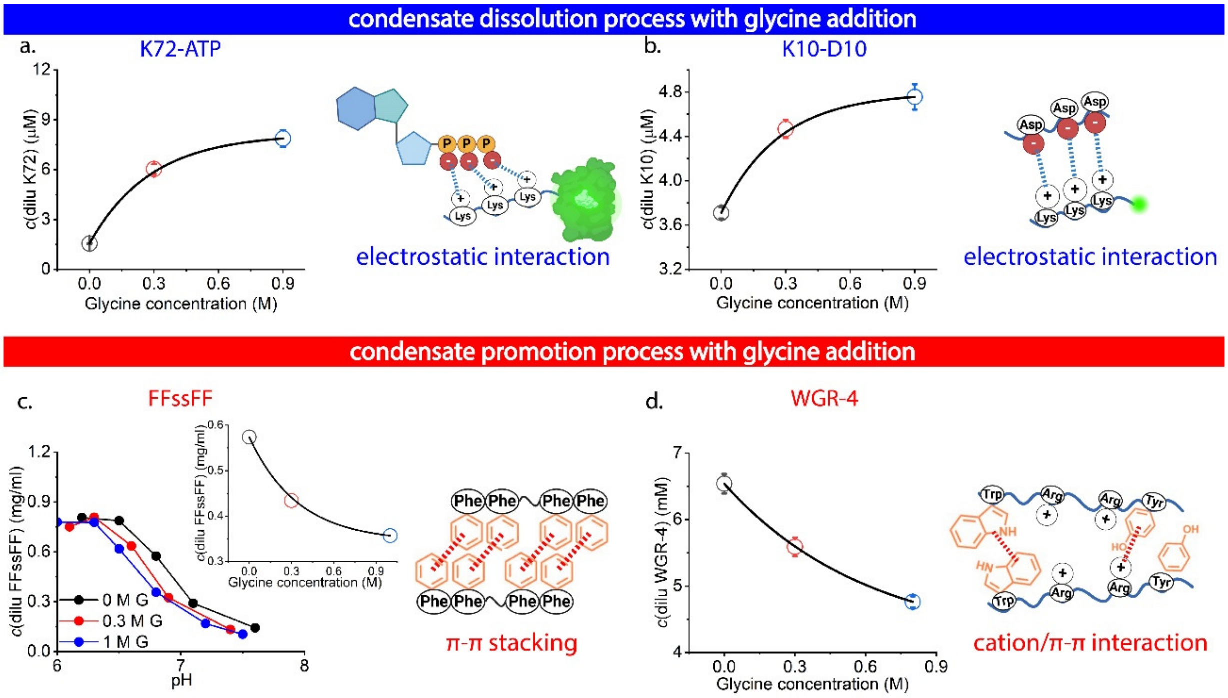


**Figure 2**: Model synthetic condensates to deconvolute the complex interaction in NPM1-RNA condensates. The peptide/protein concentration in the dilute phase after the addition of glycine (0, 0.3, and 0.9 M) as well as the expected intermolecular interactions underlying the condensate formation of a. K72-ATP system; b. K10-D10 system; c. FFssFF system (the inlet shows the FFssFF concentration in the dilute phase at pH 6.8) and d. WGR-4 system.

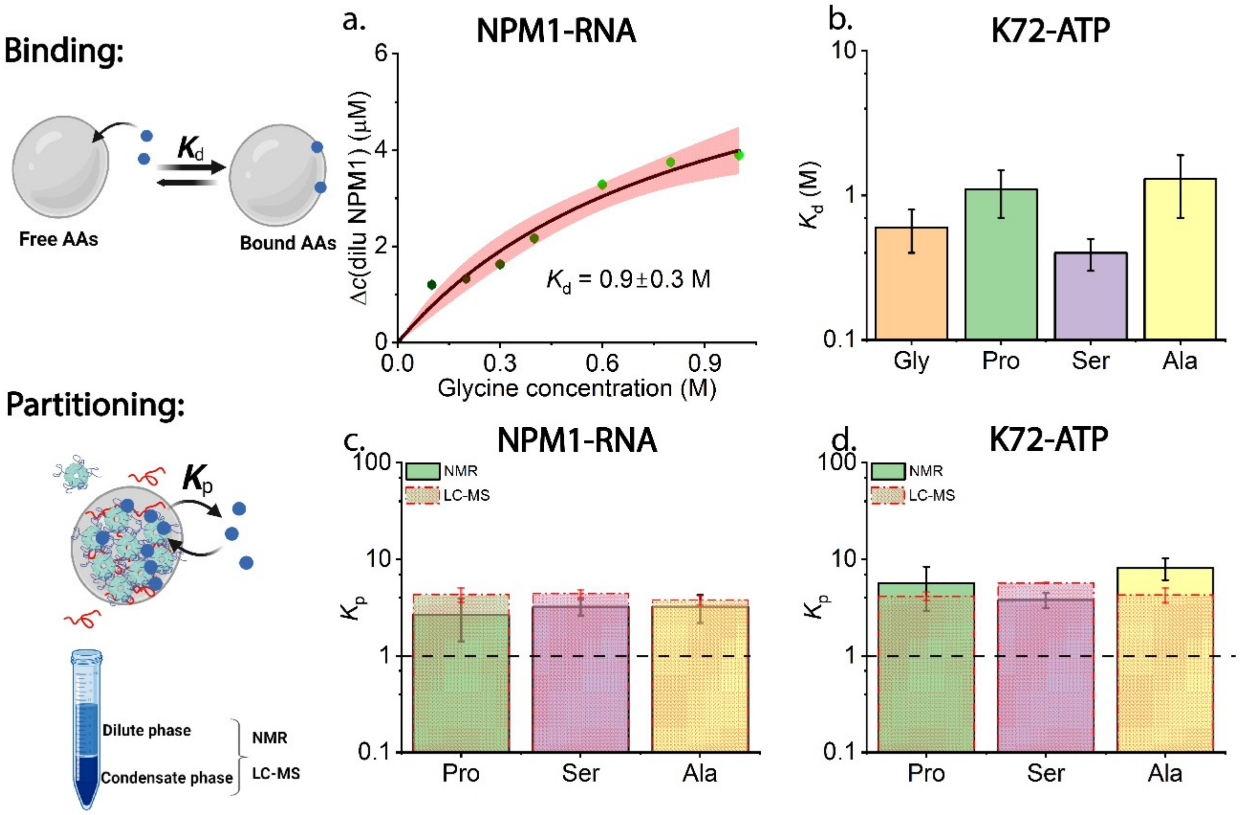


**Figure 3**: Binding and partitioning of AAs in NPM1-RNA and K72-ATP condensates. a. The NPM1 protein concentration in the dilute phase after adding glycine at varying concentrations and the fitting curve (black) with 95% confidence band (red) using the simple Langmuir-type binding model; b. The fitted binding affinities ($K_d$) for four different AAs (glycine, proline, serine, and alanine) on K72-ATP condensates; The measured partition coefficients ($K_p$) by both NMR and LC-MS for three representative AAs (proline, serine, and alanine) in the condensate phase of c. NPM1-RNA systems and d. K72-ATP systems.

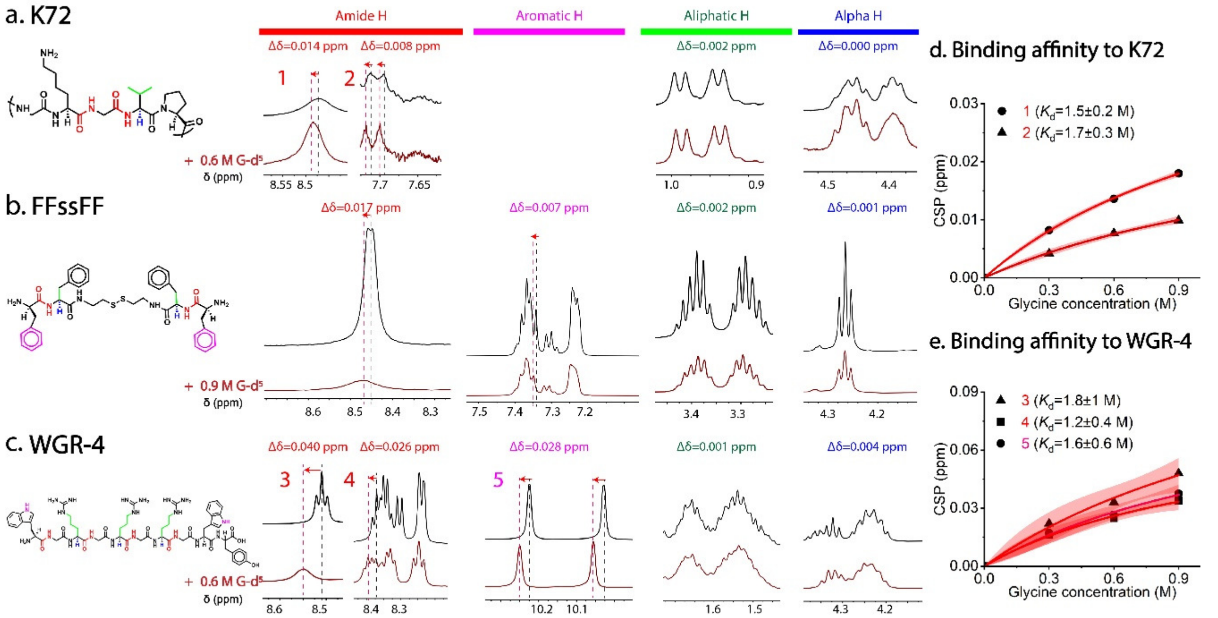


**Figure 4**: Binding of glycine (G-$d_5$) to different proteins/peptides from $^1$H NMR spectroscopy. The chemical structures, the proton peaks that experience significant chemical shifts (in red and pink) and lack significant chemical shift perturbations (in green and blue) with (red spectrum) and without (black spectrum) G-$d_5$ for a. K72; b. FFssFF and c. WGR-4; The chemical shift perturbation (CSP) of d. K72 and e. WGR-4 with the titration of G-$d_5$ at 4 different concentrations to estimate the binding affinity ($K_d$) of G to K72 and WGR-4. The fitting curves are in black with 95% confidence band (red) using the simple Langmuir-type binding model.

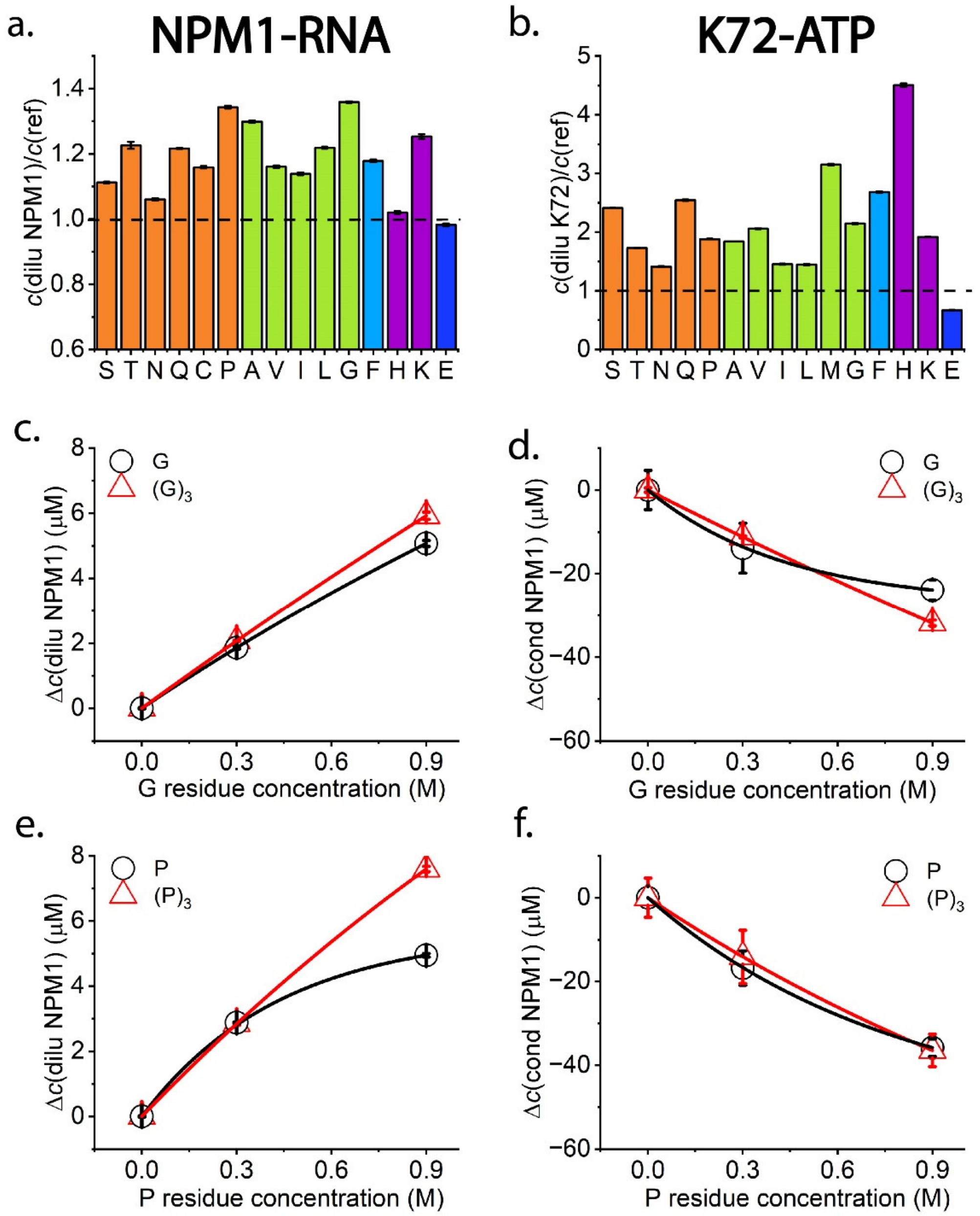


**Figure 5**: General modulation effects of AAs and the transferability to short peptides. a. NPM1 concentration in the dilute phase after the addition of different AAs (200 mM of S, T, Q, C, G, P, A, V; 100 mM of H, N, I, L, F; 20 mM K, E) divided by reference NPM1 concentration in the dilute phase in the absence of any AAs; b. K72 concentration in the dilute phase after the addition of different AAs (200 mM of S, T, Q, G, P, A, V; 100 mM of M, H, N, I, L, F; 20 mM K, E) divided by reference K72 concentration in the dilute phase in the absence of any AA; AAs are groups in colours by side chain properties. Orange, green blue and purple bars represent polar uncharged, nonpolar aliphatic, nonpolar aromatic and slightly positive charged side groups respectively; c-d. NPM1 concentration changes in the dilute and condensate phase after the addition of G and $(G)_3$ at the same ionic strength; e-f. NPM1 concentration changes in the dilute and condensate phases after the addition of P and $(P)_3$ at the same ionic strength.

In addition, we employed synthetic condensates formed by the electrostatic interaction between poly(diallyldimethyl-ammonium chloride) (PDDA) and poly(acrylic acid) (PAA), that do not contain amide groups in their backbones. We found that the addition of glycine did not affect the phase separation of this system (**Figure S17**). This experiment further supports our hypothesis that AAs bind to amide groups in the backbone of disordered proteins and modulate the protein backbone-backbone interactions, which contribute to condensate stability. AAs do not suppress phase separation when there are no amide groups in their backbone to bind to, such as in the case of PDDA-PAA condensates. In addition, the binding of AAs to side-chain aromatic groups can increase the polarity around aromatic rings, which may enhance cation/π-π interaction[25,26,40]. This explains that AAs can promote the formation of aromatic-rich FFssFF and WGR-4 condensates.

According to our proposed molecular mechanism, no specific AA side chains are needed for the observed interaction with BC components. This is supported by our findings that most AAs have similar modulation effects (**Figure 5**). More importantly, the observed transferability of the modulation to short peptides indicates that short peptides can bind reasonably strongly to disordered protein backbones, and opens the way for the rational design of short peptides to modulate BCs by strategically engineering side-chain interactions at specific binding sites. We can harness this biocompatible and versatile peptide platform to specifically target the properties and functions of physiologically important biomolecular condensates in the future[41–43]. These strategies may help to control the biological processes related to BCs, and potentially suppress undesired aging of condensates, which has been linked to a variety of neurodegenerative diseases[8,44].

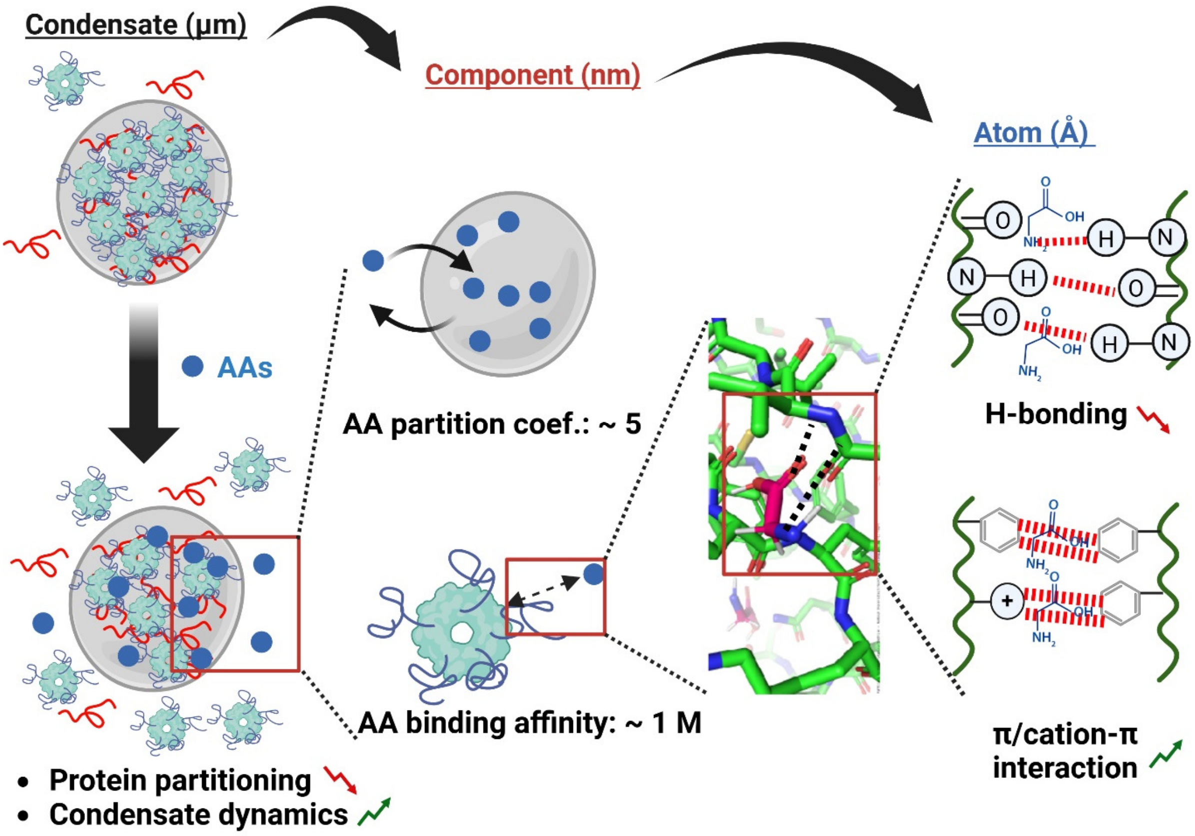


**Figure 6**: The proposed multiscale mechanism for the modulation effect of AAs on BCs, ranging from μm-scale condensate formation and dynamics, nm-scale component interaction and partitioning to Å-scale atomic interaction. Created by Biorender.

Diffusion coefficients measured by Raster image correlation spectroscopy (RICS)

The Raster Image Correlation Spectroscopy (RICS) was performed on a Leica SP8 confocal microscope equipped with a single-photon detector. Calibration of the focal volume waist $\omega_0$ was performed using the known diffusion coefficient of Alexa 488 of 435 $\mu m^2/s$ (T = 22.5 ± 0.5 °C) in water, and $\omega_z$ was set to 3 times the value of $\omega_0$[46]. All measurements were captured at a resolution of 256 × 256 pixels with a 20 nm pixel size using a 63x oil objective. Condensates were measured at 10 Hz line speed with 15 frames acquired per data point. Analysis of autocorrelation curves was done using PAM[47].

Langmuir-type binding model fitting of dilute phase NPM1 concentration change and the chemical shift perturbation with different glycine concentrations

The dilute phase NPM1 concentration change and the chemical shift perturbation (both denoted as $\Delta$) with different AA concentrations ($c$) was fitted by a simple binding model[15,36] under the assumption of excess AA: $\Delta = \frac{\Delta_{max} \times c}{K_d + c}$.
